# Mitochondrial DNA haplotypes reveal a fine-scale population structure in sperm whales (*Physeter macrocephalus*) around São Miguel Island (Azores)

**DOI:** 10.64898/2026.07.21.739783

**Authors:** Stéphanie R.A. Suciu, José M.N. Azevedo, Seán A. O’Callaghan, Bruno Serranito, Jean-Luc Jung

**Author notes:** **Correspondence:** Stéphanie Suciu.

## Abstract

The sperm whale is an emblematic species of the Azores archipelago. After a century of intensive whaling, the species has become a focal point for ecotourism since the late 1980s, enabling collaborative research during tourism activities. This study assessed the feasibility of using a citizen science based non-invasive sampling approach to evaluate the genetic diversity of local sperm whales. Seven local whale-watching companies contributed to collect 70 fecal samples and 34 sloughed skin fragments between 2019 and 2024. A 588-bp fragment of mitochondrial DNA Control Region (MCR) was successfully sequenced from 56 fecal samples and all skin fragments, yielding 90 MCR sequences assigned to 79 individual sperm whales. MCR sequences from both sources of the same individuals were in perfect agreement, validating the use of fecal samples for mitochondrial haplotype analysis - the first study to do so in sperm whales. Four MCR haplotypes, including a newly determined one, evidenced different maternal lineages within the Macaronesia population. Haplotypes showed a clear geographic pattern around São Miguel Island: one was restricted almost exclusively to the south coast, while the other three occurred predominantly on the north. This contrasted distribution is likely correlated to different matrilineal groups.

## 1. Introduction

Sperm whale society is structured around stable matrilineal social units, typically composed of related adult females and their offspring. These units represent the core of social organization of the species (Eguiguren et al., 2023; Gero et al., 2013; Sarano et al., 2021; Whitehead & Kahn, 1992). They may be composed of 3 to over 20 animals (Cantor et al., 2019; Konrad et al., 2018; Sarano et al., 2022; Whitehead et al., 1991) and can, sometimes, include unrelated individuals (Konrad et al., 2018; Sarano et al., 2021). Together, they cooperate in calf care (Sarano et al., 2023), communal defense against predators like killer whales (*Orcinus orca*) (Pitman et al., 2001), foraging and knowledge sharing (Gero et al., 2013; Sarano et al., 2023; Whitehead et al., 1989; Whitehead, 2003), and often travel together, maintaining long-term associations within warm-temperate and tropical waters, often not exceeding 45°N (Mullin et al., 2022; Whitehead, 2018). The movement of matrilineal groups is characterized by regional fidelity, displaying coordinated, likely democratic, group decisions about travel (Whitehead, 2016), with groups covering significant distances together each day (Konrad et al., 2018; Vachon et al., 2022), adapting to the distribution of resources (Mizroch & Rice, 2013; Whitehead et al., 2008).

These social and geographical fidelities result in genetic differentiation between ocean basins (Alexander et al., 2016; Ferreira et al., 2022; Lyrholm & Gyllensten, 1998; Whitehead et al., 1992), but also at smaller scales, within semi-enclosed seas. For instance, studies have shown distinct mitochondrial DNA haplotypes for females from the Caribbean (Gero et al., 2014), as well in the Mediterranean (Engelhaupt et al., 2009), compared to the rest of the North Atlantic. Sperm whale social units may regularly associate with other units, leading to temporary groups that can last from hours to days, yet they do not group with all units present in the same area (Cantor et al., 2019; Whitehead et al., 2012), indicating selective social preferences consistent with clan structures defined by vocal dialects (Christal et al., 1998; Gero et al., 2014; Rendell et al., 2012; Whitehead, 2024). Moreover, Vachon et al. (2022) showed that social structure may drive differential space use, highlighting the need to incorporate social clans into population management and conservation frameworks (Engelhaupt et al., 2009; Lizewski et al., 2025; Whitehead, 2016).

Males, in contrast, leave their natal units during adolescence, forming loose bachelor groups and with solitarily periods at higher latitudes (Best 1979; Morange et al, 2026; Whitehead 2003). They only return periodically to warmer waters for breeding in social groups, likely distinct from the one of their births (Lydersen et al., 2025; Mendes et al., 2007; Morange et al., 2025; Steiner et al., 2012, Girardet et al., 2022) facilitating genetic mixing (Mesnick et al., 2011), although genetic evidence for such patterns remains lacking in the Atlantic. In the Indian Ocean, males appear to exhibit a degree of social fidelity to matrilinear group, returning to visit them over time (Girardet et al., 2022).

Sperm whales are one of the emblematic marine species of the Azores. Whale watching occurs off five of the nine islands of the Azores archipelago (Figure 1). São Miguel, the largest of these islands, is the most popular destination for tourists (Serviço Regional de Estatística dos Açores, 2023; Observatório do turismo dos Açores). In contrast, Pico, Faial, São Jorge, and Terceira are noted for having a higher frequency of cetacean sightings (Silva et al., 2003). In the late 1980s, the shift from whaling to whale watching created new avenues for marine research (Ellis, 2009; Oliveira et al., 2007) by offering the possibility of a participatory research approach. Whale watching crews, often composed of staff with an academic background (biologists), collect valuable ecological data from their daily observations, thus providing a cost-effective alternative to scientific data collection (Coché et al., 2021; García et al., 2023; Suciu et al., 2025) and complementing traditional scientific surveys, particularly in regions lacking baseline data or where funding is limited. They can also participate in sample collection campaigns, particularly those that rely on non-invasive sampling methods, such as collecting sloughed skin and fecal samples. Such samples can significantly advance the study of sperm whale ecology and genetics (e.g. analyses of population structure, kinship, individual identification, (Byrne et al., 2021). These approaches can strongly help effectively and, more importantly, ethically monitor elusive marine species (Amos et al., 1992; Girardet et al., 2022; Sarano et al., 2021).

**Figure 1.**
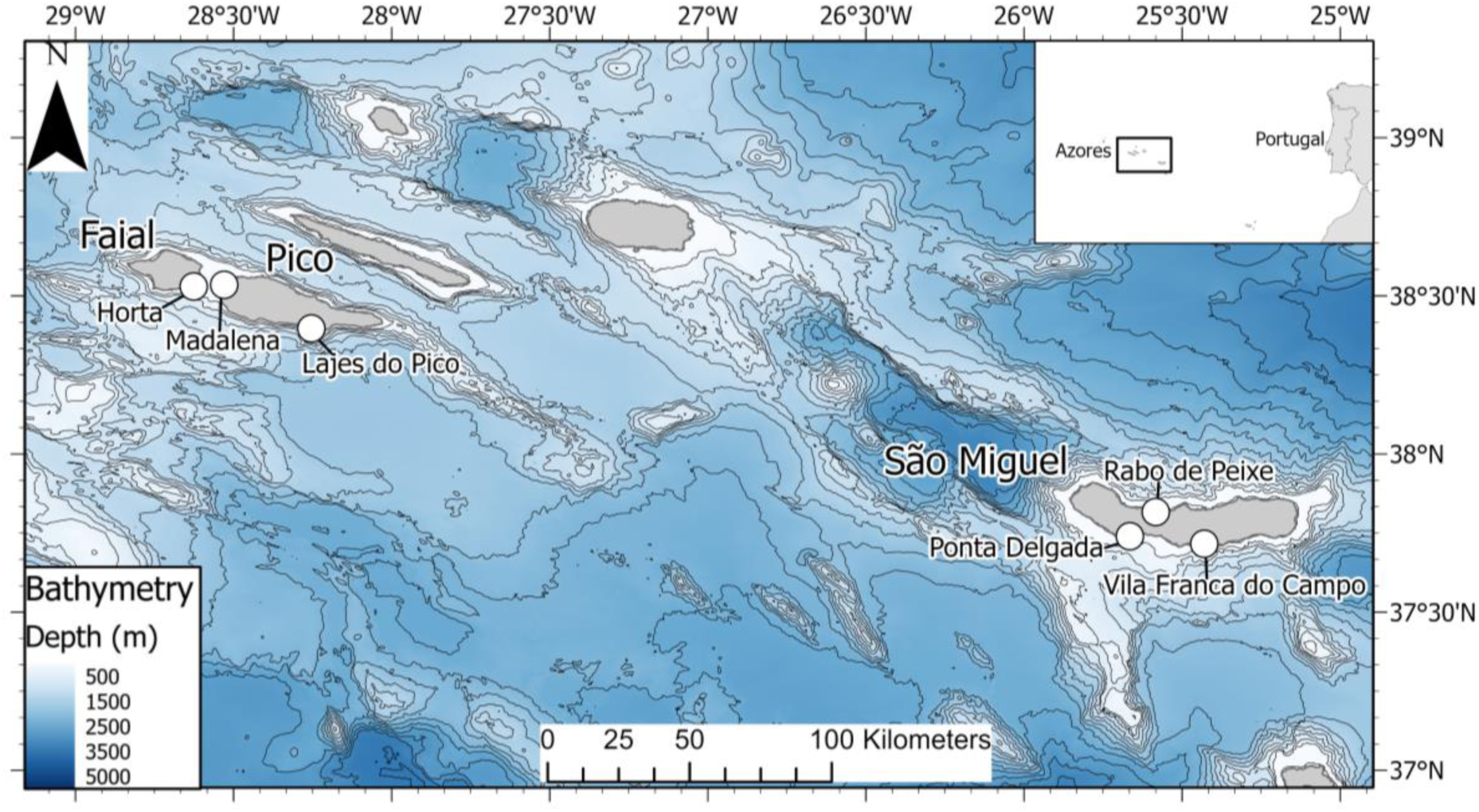
Map of the study area, highlighting São Miguel Island, Faial and Pico Islands on the archipelago of the Azores in the central and eastern groups, and location of the main harbors where boats departed from.

Most of what we know about sperm whale social structure is based on research in the Indian (Sarano et al., 2021), Pacific (Whitehead, 2003) and Atlantic basins (Brennan et al., 2026; Cantor et al., 2019; Gero et al., 2015). Azorean sperm whales form part of the larger population of Macaronesia (Boys et al., 2019; Pinela et al., 2009), with a subset of females that returns annually to specific archipelagos, whereas others use the area transiently (van der Linde and Eriksson 2020; Ferreira et al., 2022). Only a few studies investigated their population and social structures (van der Linde & Eriksson, 2020). A study based on photo-identification has determined 12 social units around São Miguel on the basis of preferred individual associations, which showed fidelity to the water around the island (van der Linde & Eriksson, 2020). Sperm whale fidelity to the archipelago or to a group of islands has been previously reported (Ferreira et al., 2022; Matthews et al., 2023; Pinela et al., 2009; van der Linde & Eriksson, 2020). However, no genetic research has yet examined fine-scale geographic preferences.

We studied the genetic diversity of sperm whales around São Miguel, using a non-invasive, citizen science-based approach by developing a dedicated protocol that involves whale watchers. We collected sloughed skin and feces samples from individuals and as often as possible identified from photographs. We then sequenced a part of the mitochondrial DNA Control region (MCR), widely used to assess genetic diversity and infer population structure and history (Ballard & Whitlock, 2004; Morin et al., 2018), validating the use of fecal samples for MCR haplotype determination. As the mitochondrial DNA (mtDNA) is maternally inherited, individuals within a matrilineal group should share the same mitochondrial haplotype (Sarano et al., 2021), enabling to explore social organization/structure of the studied individuals.

We studied sperm whales primarily around São Miguel, testing the null hypothesis that mitochondrial haplotype repartition is not correlated to their spatial distribution around the island. A non-random distribution would suggest that social units harboring different haplotypes either (1) exhibit spatial segregation (e.g. avoidance or cohesion with haplotypes) and/or (2) display distinct ecological patterns.

## 2. Material and methods

### 2.1. Study area and sampling campaign

Sperm whales are regularly observed in the Azores (Portugal) and represent a focal species for whale watching companies. To maximize sampling effort and resources, this study combined a participatory science approach involving seven whale watching companies and dedicated research trips. The main investigator was based in São Miguel, facilitating direct collaboration with whale watching operators.

The rigid inflatable boats (RIB) from whale watching companies departed from the harbors of Ponta Delgada, Vila Franca do Campo and Rabo de Peixe (São Miguel Island), from Madalena and Lajes do Pico (Pico Island) and from Horta (Faial Island). The dedicated research boats, a fiberglass boat (<12m) or RIB’s were based on São Miguel Island, departing from Vila Franca do Campo, Ponta Delgada and Rabo de Peixe (Figure 1). The study area was located up to 25 km off the coast depending on where whales were located. Boats were directed from a land-based lookout (vigia) to the location of whales when an observer was available, and a directional hydrophone was also used to locate and acoustically track animals when needed. Samples were collected between 2019 and 2024, with a peak during the high tourist season from May to September, corresponding to better weather conditions and increased seasonal tourism.

Most of the samples were collected around the island of São Miguel (n = 93), with a small proportion (n = 11) coming from the central group of islands, around Pico and Faial (Figure 1).

### 2.2. Collection of samples and DNA extraction

Sperm whales were individually identified using natural marking on their flukes/dorsal fins (Arnbom T, 1987; van der Linde & Eriksson, 2020) from photos collected with handheld digital cameras or from overhead using a drone, under the required permits (O’Callaghan et al., 2024). The procedures from the applicable legislation for cetacean observation (Decreto Legislativo Regional n° 9/99/A de 22 de Março Decreto Legislativo Regional n° 10/2003/A and Portaria 5/2004) were strictly followed. Fieldwork was carried out under Azores Government licences.

After a sperm whale fluked up when going on a deep dive, the boat slowly approached the fluke print (glassy area of water caused by the tail upwelling) which was visually scanned for any biological material (fecal matter and sloughed skin) (Figure 2a).

**Figure 2.**
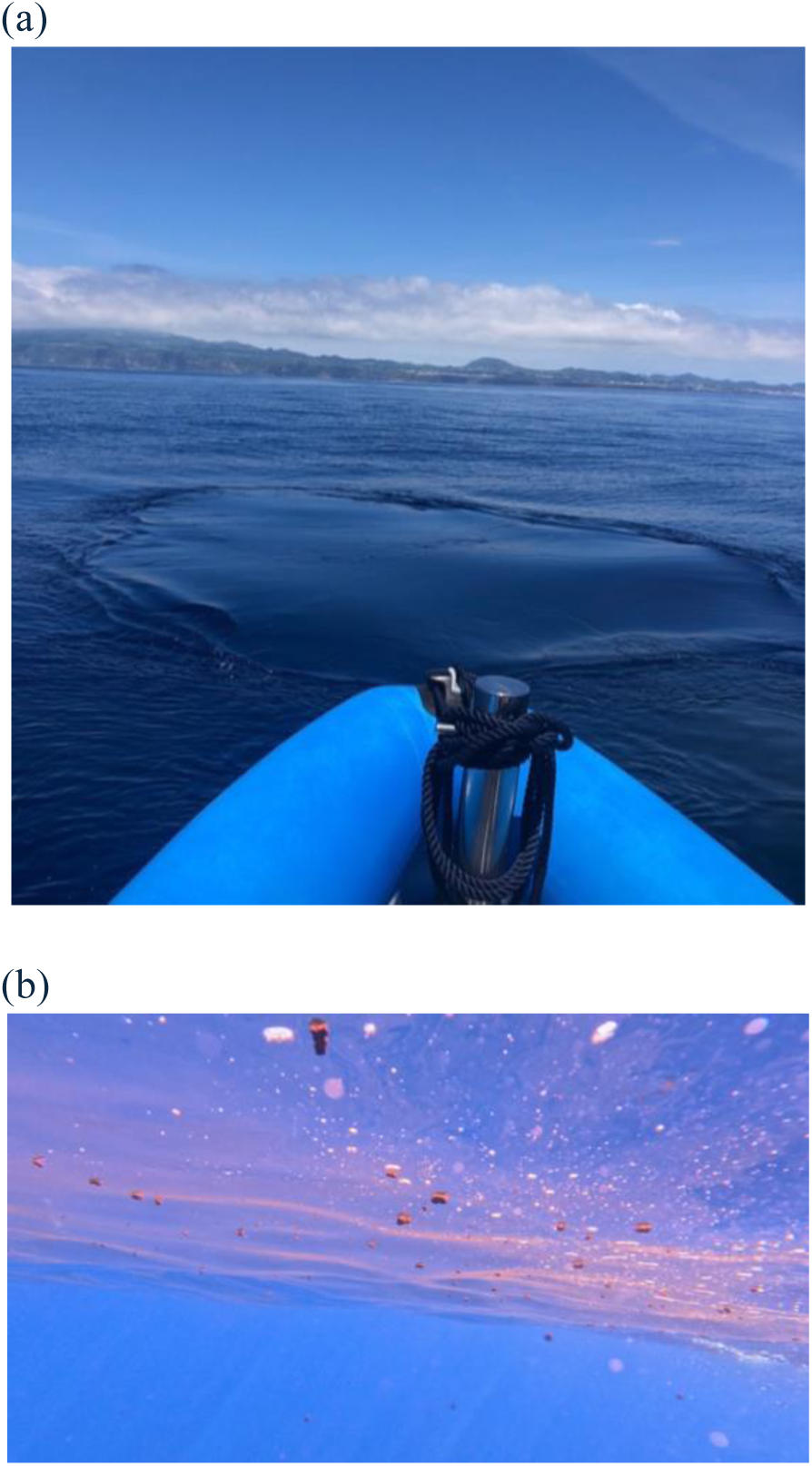
Fecal sample collection (a) Approaching a sperm whale’s fluke print after its feeding dive. Note that the fluke print is clearly visible as a more or less circular area of glassy water (b) Sperm whale fecal matter just under the surface. Photos by Stéphanie R.A. Suciu.

Sperm whales regularly defecate before diving, producing feces that range from dispersed plume to floating semi-solid clumps (Figure 2b). Fecal samples were collected using two different protocols: subsamples “F” were collected when feces were found in a high concentration or as clumps, directly in a sterile 100mL polypropylene collection pot or with a fine mesh net, then preserved in 96% ethanol or kept on ice, and stored at -20° C; for subsamples “E”, seawater containing the fecal material was filtered with single-use environmental DNA (eDNA) dual-filter capsules with 0.8 µm PES membranes (Sylphium molecular ecology, Groningen, The Netherlands) following manufacturer instructions, then stored at 4°C. These subsamples were labeled with an ‘E’ or ‘F’ identifier and sequential number. When possible, both protocols were used so, both types of fecal subsamples were collected from the same sperm whale. Equipment was carefully rinsed with tap water and bleached to prevent cross-contamination.

Skin samples of sloughed epidermis were found at the surface, released as the act of surfacing and diving repeatedly allowing skin to shed. Occasionally, skin samples were also found after social interactions where animals are in close contact, hampering efforts to determine the individual the sample originated from. Skin fragments were recovered with a bucket and transferred to a 50mL falcon tube. Sea water was replaced by 96% ethanol, and samples were held at room temperature until return and stored at 4° C and labelled with an ‘S’ identifier and sequential number. When possible, both types of samples (fecal and skin) were taken on the same sperm whale.

The DNA from fecal samples was extracted according to the subsamples type: for subsample “F”, the DNeasy Blood & Tissue Kit (Qiagen, Hilden, Germany) was used, following the manufacturer’s instructions; for subsample “E”, the DNA Isolation Kit SYL002 (Sylphium Molecular Ecology, Groningen, The Netherlands) was used, with the protocol adapted to process 1 ml of eDNA sample in CTAB buffer per extraction. The DNA from skin samples was extracted using the PureLink Genomic DNA Mini Kit (Invitrogen, Waltham, MA, USA), according to the manufacturer’s guidelines. In case of undetected PCR products (see part 2.4), the genomic DNA extract was subjected to a cleanup step using the NucleoSpin Inhibitor Removal Kit (Macherey-Nagel, Düren, Germany) to remove potential PCR inhibitors, following the manufacturer’s instructions.

The concentration and purity of the extracted DNA were measured spectrophotometrically using a NanoDrop 100 (Thermo Fisher Scientific, Waltham, USA). DNA extracts were stored at -20°C. Unsuccessful DNA extraction and PCR reactions were repeated up to three times.

### 2.3. PCR amplification and sequencing of mtDNA control region

The DLP1.5 and DLP8G primers (Garrigue et al., 2004) were used to amplify a 774bp fragment of the mtDNA control region (MCR). The PCR products were analyzed by electrophoresis on a 2% (p/v) agarose gel. Amplicons of the right size were purified and sequenced by external providers (Stabvida, Caparica, Portugal; Eurofins, Germany) using an ABI 3730XL DNA Analyzer (Applied Biosystems). For each feces or skin sample, at least one PCR product was sequenced in both directions using Dlp1.5 and Dlp8G primers (Garrigue et al, 2004). All sequences were checked for quality using Geneious Prime v11.0 (Biomatters; Kearse et al., 2012).

### 2.4. Analysis of MCRs polymorphisms and determination of haplotypes

Sequences were aligned with Geneious Prime v11.0. A common 588bp MCR sequence region was determined unambiguously from all samples, corresponding to nt 15516 to 16104 of the 16429 bp complete mitochondrial genome (NCBI accession number KU891354.1, Morin et al., 2018) and including the hypervariable region of the D-loop (Alexander et al., 2012). Haplotypes were identified using the DNAsp 6 software (Rozas et al., 2017) and the final haplotype sequences were compared with those available in GenBank using BLAST (Altschul et al., 1997). They overlapped with the 399bp sequence from Mesnick et al, (2011), defining MCR haplotypes: A (Accession No. DQ512921), B (DQ51292) and C (DQ512923).

To compare the genetic diversity of the Azorean sperm whales with all individuals studied from the North Atlantic, a new dataset was created, grouping individual sequences: 79 of this study (see 3.1), 45 from Lyrholm & Gyllensten, 1998, 102 from Engelhaupt et al., 2009 and 31 from Morin et al., 2018 (Table S2). After alignment, a common overlapping sequence of 359bp was determined, and no CR haplotype from the Azores was lost in this process. Individuals from the Gulf of Mexico and the Mediterranean were not included in the North Atlantic comparison as these populations showed a particular haplotype tendency that is different from the rest of the North Atlantic (Engelhaupt et al., 2009; Ortega-Ortiz et al., 2012).

To assess the molecular diversity, we used DNAsp 6 software (Rozas et al., 2017) to calculate the number of haplotypes (Nh), the number of polymorphic sites (k), haplotype diversity (*Hd*) and nucleotide diversity (*π*). DNAsp was also used to perform neutrality tests; we estimated Fu’s *Fs* (Fu, 1996) and Tajima’s *D* (Tajima, 1989) to assess changes in population demography, such as expansion or bottleneck.

A median-joining network was constructed with Network software (Bandelt et al., 1999) to infer the relationships and the number of substitutions between haplotypes graphically.

### 2.5. Evaluation of mitochondrial DNA spatio-temporal variations and correlation with possible differences in habitat

To investigate whether the individuals showed a local geographical distribution correlated to their mitochondrial haplotypes, samples were divided into three geographic groups according to their sampling location around São Miguel: south, north (Figure 3) and an additional central group. Genetic differentiation was then tested among geographical groups using a χ^2^ test based on haplotype frequencies (Hudson et al., 1992) and a nearest neighbor statistics (*S_NN_*) index (Hudson, 2000), which was suitable for evaluating population structure when sample sizes are small. Statistical significance was assessed via 1,000 random permutations. Molecular and haplotype differentiations were estimated with, respectively, *Φ_ST_* and *F_ST_* (Hudson et al., 1992), using Arlequin 3.5.1 (Excoffier & Lischer, 2010). These same indices were calculated between the samples collected in 2022, 2023 and 2024 to assess possible temporal differences (the other years were excluded due to insufficient sample sizes).

**Figure 3.**
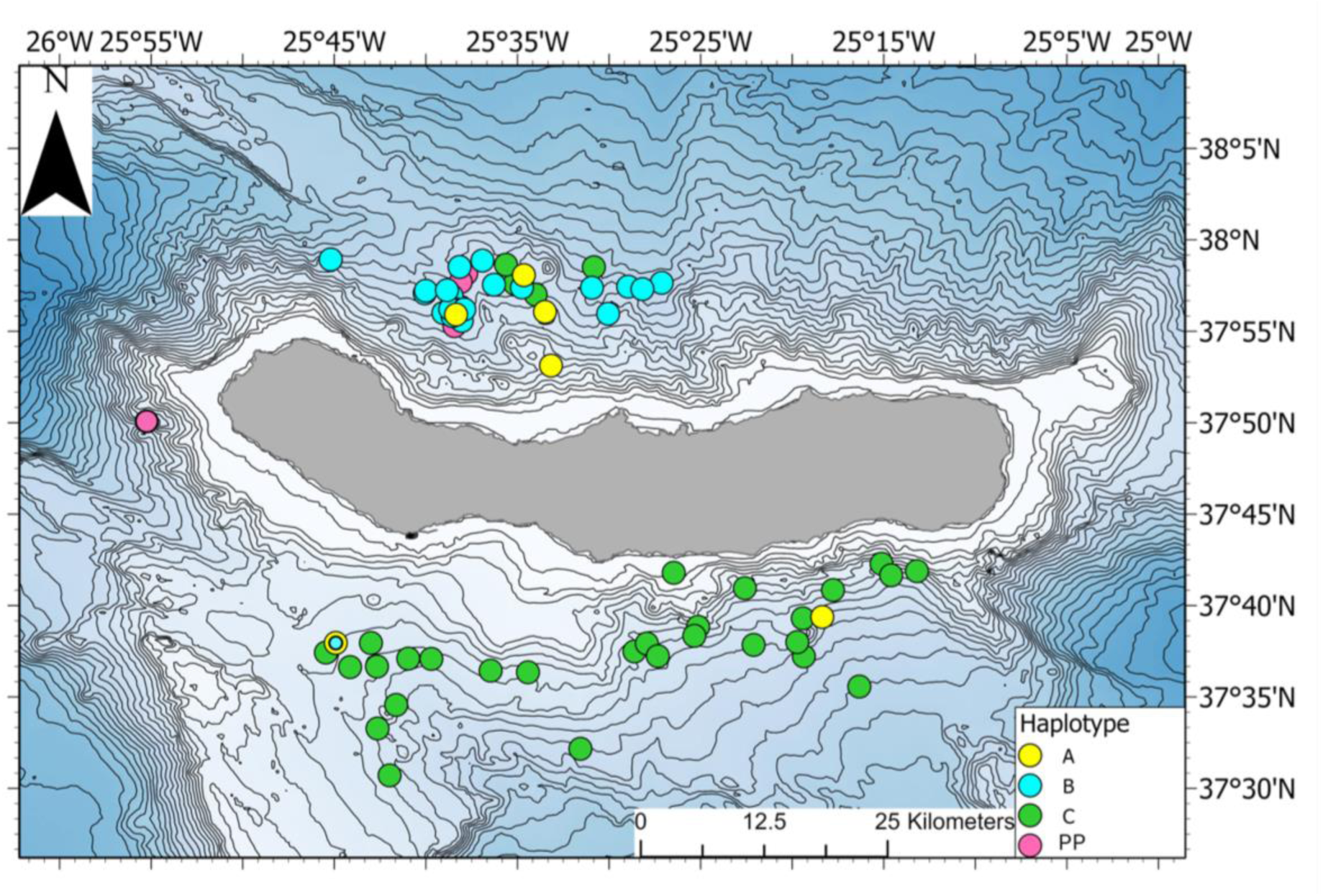
Distribution of the haplotypes on the studied site of São Miguel (all the samples for which geographical data were available (n= 72) are represented).

To investigate whether foraging depth preferences differed among sperm whales with different mitochondrial haplotypes, we used a harmonized Digital Terrain Model (DTM) from the European Marine Observation and Data network (EMODnet) with a resolution of 1/16 arc minutes (∼115 m) (EMODnet Bathymetry Consortium, 2024). For each individual, we extracted the bathymetric value at the coordinates of the last surface observation preceding a deep dive, used as a proxy for feeding depth (Drouot et al., 2005). That value was assumed to reflect local foraging conditions, as depth is a known driver of cephalopod distribution (Rosa, 2013). Depth variation between haplotypes was assessed using a Linear Mixed Model (LMM) as implemented in the R package *glmmTMB* (Brooks et al., 2017) with haplotype as fixed effect and using individual ID, year and month of the observation as random effects to account for potential temporal and individual-dependant variability. We checked model assumptions of normality in residuals and by visually inspecting QQplots using the R package DHARMa (Hartig, 2016) (Table S4, Figure S1). Post-hoc pairwise comparisons were performed using the R package *emmeans* (Lenth et al., 2024) to evaluate the differences in depth between haplotypes after accounting for covariate effects (Table S5).

## 3. Results

Over 52 days between 2019 and 2024, we collected 104 samples from up to 91 individuals, identified as often as possible (Table S1). There were 70 fecal samples from 66 individual whales and 34 skin samples from 31 individual whales, including six individuals sampled for both fecal and skin samples (Table S1). Indeed, some animals were resighted (n=2), usually on the same day resurfacing after a deep dive within the same area (PmM64, AZ29), while others (n=2) were resighted several months/years apart (PmM76, Mr Liable). For the mature male, nicknamed “Mr. Liable, six samples were collected on five different days over three years (2019, 2022, 2023). In some cases, both skin and fecal samples were found on the same fluke print (n = 4). These individuals sampled several times allowed to validate the methodology (see 3.1). Besides the sperm whales re-sampled several times, other samples come from different individuals. The details of the samples collected per individual are given in Table S3. For individuals encountered on different days, without photo-identification available (n=58), we assumed that none were re-encountered. This decision was made based on aerial photo-identification that showed no recaptured individuals between 2022 and 2024 on the north coast of São Miguel (O’Callaghan et al., 2024).

All fecal samples were found and collected on fluke prints, ensuring minimal environmental contamination and maximizing sample integrity: 18 were collected following the protocol for subsample “E”, 23 following the protocol for subsample “F”, and 29 both types of protocols (“E” and “F”) were successfully gathered. From the 34 sloughed skin collected: 28 were found on a fluke print, 5 next to animals during social behavior, and one was from a stranded juvenile that came ashore on the South of São Miguel in January 2025 (Figure S1).

### 3.1. Success and reproducibility of MCR sequencing

The analysis of the 104 samples yielded 90 MCR sequences covering the 588bp MCR fragment, demonstrating a high amplification success across the two types of non-invasive samples. Sequencing success reached 80% for fecal samples, and 100% for the sloughed skin samples (Table1, Figure S2).

Comparisons of the MCR sequences obtained from different samples taken from the same animals (two fecal protocols, “F” and “E”, in Table S6; fecal and skin samples in Table S7) were all similar regardless of protocol or sample type.

Overall, the 90 MCR sequence determined were assigned to 79 different individuals, of which 33 were photo-identified.

### 3.2. MCR polymorphisms and haplotype characterization

We identified four different haplotypes in our dataset, each separated from the others by a single mutation (Table S8). Three of the four had a 100% similarity on their common part with, respectively, the sequences of haplotypes A, B and C defined by Mesnick et al, 2011 (Table S9). We therefore named them as SMG for São Miguel as most of our samples came from around this island, following the Mesnick et al (2005) nomenclature: SMG_A, SMG_B, SMG_C (Accession no: PZ311571 - PZ311573). One haplotype was not previously described in the literature. This new haplotype was named SMG_PP (Accession no: PZ311574), as “PP” is the next logical name based on the initial given by Alexander et al, 2016 (Table S9).

The 79 individual MCR sequences enabled the calculation of haplotype (*Hd*) and nucleotide (π) diversities (Table 3). Haplotype diversity was higher in the samples taken off the North of São Miguel and in the central group, than on the South of São Miguel, which also exhibited significantly lower nucleotide diversity than the other groups (Table 1). Values for Tajima’s *D* and Fu’s *Fs* were positive, but not statistically significant.

**Table 2.** Sample types and success rates of MCR sequencing per type of sample.

| Type of sample |  | Number of samples | Number of analyzed samples | Number of samples with successful MCR determination (success rate) |
| --- | --- | --- | --- | --- |
| Feces | Total | 70 | 70 | 56 (80%) |
|  | Subsample “F” | 52 | 46 | 34 (74%) |
|  | Subsample “E” | 47 | 46 | 35 (76%) |
| Skin |  | 34 | 34 | 34 (100%) |

**Table 3.** mtDNA control region haplotype diversity statistics of the group of samples. *n: number of individuals, l: sequence length (bp), K: average number of differences; h: no. of haplotypes, Hd: haplotype diversity, π: nucleotide diversity; SMG: São Miguel; central group: Faial and Pico. n.s: non-significant values*

| | n | l | h | $Hd$ | $\pi$ (%) | Tajima’s $D$ | Fu’s $F_s$ |
| --- | --- | --- | --- | --- | --- | --- | --- |
| SMG South | 36 | 588 | 4 | 0.303 | 0.077 | - | - |
| SMG North | 35 | 588 | 4 | 0.629 | 0.128 | - | - |
| Central group | 8 | 588 | 3 | 0.607 | 0.134 | - | - |
| Total | 79 | 588 | 4 | 0.674 | 0.155 | 0.931(ns) | 0.989(ns) |

**Table 4.** Pairwise genetic differentiation (F_ST_) among spatially defined populations.

| Fst | n | São Miguel North | São Miguel South | Central group |
| --- | --- | --- | --- | --- |
| São Miguel North | 35 | 0 | - | - |
| São Miguel South | 36 | 0.435* | 0 | - |
| Central group | 8 | 0.192* | 0.531* | 0 |

\*p-value < 0.0001 (Number of permutations: 110)

### 3.3. Geographical segregation per haplotype and foraging depth differences

The spatial distribution of sperm whale haplotypes around São Miguel Island showed a clear geographic pattern (Figure 3): Haplotype SMG_C was almost totally restricted to sperm whales sampled on the south of São Miguel, whereas haplotypes SMG_A, SMG_B and the newly identified haplotype SMG_PP are mainly found in the north of the island. Individuals carrying the SMG_PP haplotype were consistently observed in a localized area on three separate days across two different years (Figure 3).

A strong genetic differentiation was therefore observed between the three geographical groups (Table 3). All indices were significant and indicate a differential geographical distribution of the haplotypes between the north and south of São Miguel, and the central group: *S_NN_* = 0.633 (p < 0.001), *χ^2^* = 54.63 (df = 6, p < 0.001), *Φ_ST_* = 0.427 and *F_ST_*_=_ 0.435.

We used feeding depth as a proxy for foraging habitat and tested whether it differed among mitochondrial haplotypes (Figure 4). The seafloor was significantly deeper for haplotypes determined on sperm whales sampled on the north coast of the island than for those sampled on the south coast, where the haplotype SMG_C was almost exclusive. On average, haplotype SMG_C were determined for samples collected at a mean depth of 962 m (SD: ±216m), whereas haplotypes SMG_A, SMG_B and SMG_PP were associated with a mean depth higher than 1,000 m (SMG_A at 1090 m (±SD 382 m), SMG_B at 1,178 m (±SD: 180 m) and SMG_PP at 1,139 m (±SD: 38 m) (Figure 4).

**Figure 4.**
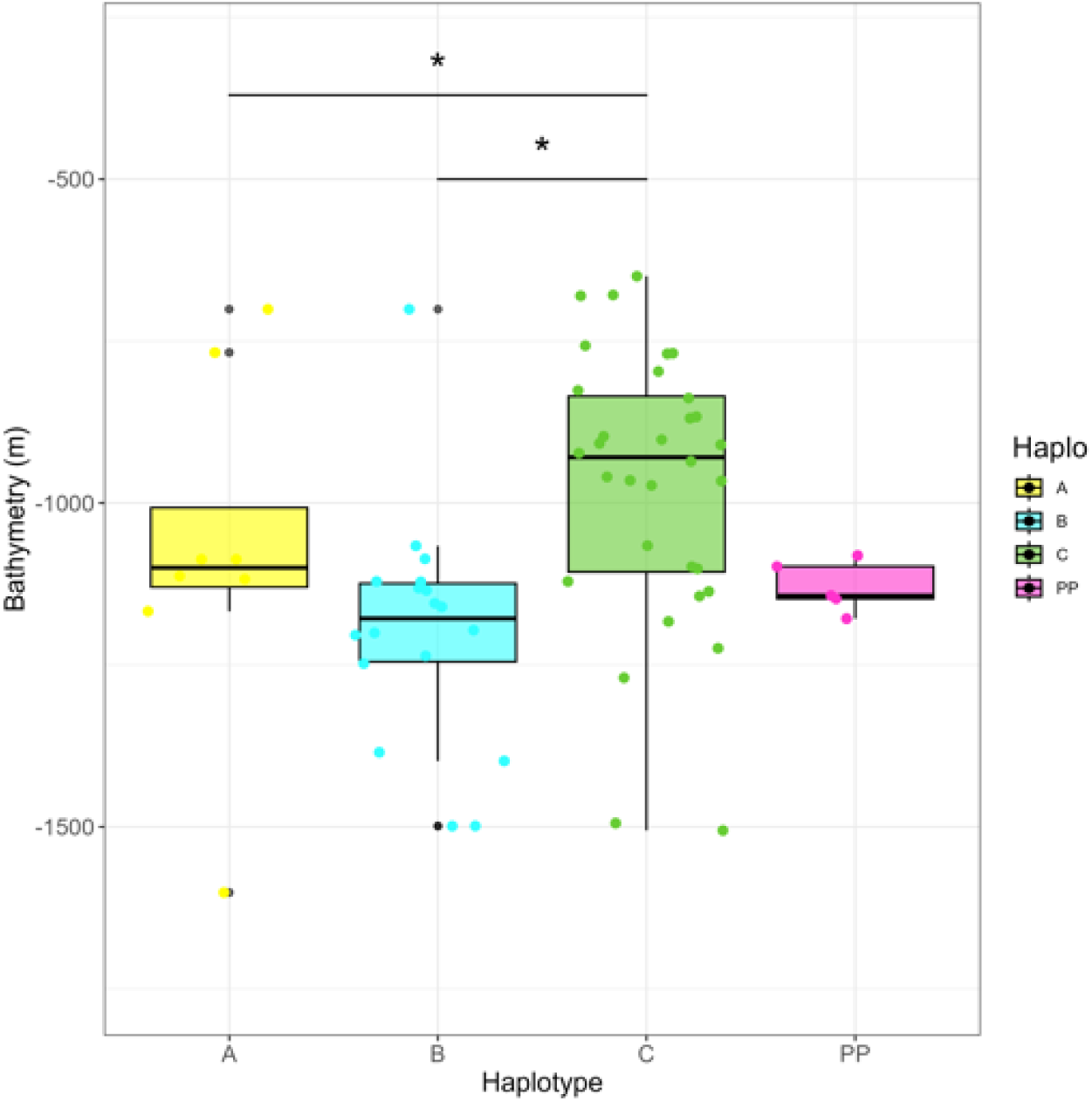
Bathymetry distribution and average depth at the sampling location per haplotypes A, B, C and PP.

### 3.4. Temporal distribution

No significant genetic differences were detected between sampling years (*χ²* = 14.414, df = 6, p > 0.1; *S_NN_* = 0.32955, PM test p-value > 0.01), which was consistent with non-significant *Φ_ST_* and *Fst* values (Table S10).

### 3.5. Comparison with North Atlantic genetic diversity

The haplotype diversity (*Hd*) and nucleotide diversity (π) for the dataset of this study were similar to the North Atlantic dataset (Table S11). Fu’s *Fs* and Tajima’s *D* indices were not significantly different.

## 4. Discussion

Recovering from the impacts of the whaling era, the sperm whale is currently classified as “Vulnerable” by the International Union for the Conservation of Nature (IUCN) (Whitehead et al., 2025). Their continued protection relies notably on a thorough understanding of the species’ genetic diversity, the spatial scales at which it is structured, and the variation in habitat use among groups. This study explored the genetic diversity of sperm whales in the Azores, focusing on their fine-scale distribution around São Miguel Island. Additionally, it implemented an ethical and inclusive approach to assess and optimize the collection of samples and the analysis of the obtained genetic data. In an economic context where funding is increasingly limited despite the urgent need to protect ocean health, participatory science and non-invasive sampling techniques coupled to innovations in molecular analysis are precious tools (Byrne et al., 2021).

### 4.1. Successful sperm whale MCR sequencing from non-invasive samples collected by participatory science

This study confirmed the feasibility of using sloughed skin to assess sperm whale mtDNA control region (MCR) haplotypes for genetic analysis in sperm whales (Girardet et al., 2022; Konrad et al., 2018; Richard et al., 1996; Sarano et al., 2021). In addition, it demonstrated for the first time the potential of also using fecal samples for sperm whales, as it has been done for other marine mammal species (Jackson, 2022; Ooi et al., 2023; Parsons et al., 1999). The success rate and consistency of MCR sequence determination was very high and confirmed by multiple samplings of the same individual whale, using different sampled material (feces and sloughed skins), and different collection methods.

This study highlighted the importance of encouraging participatory science in addition to dedicated scientific research campaigns to study marine biodiversity in general, and to collect sperm whale non-invasive samples in particular, using fit for purpose and standardized sampling protocols. Half of the fecal and sloughed skin samples of this study were collected during touristic activities by seven different companies across three islands. Engaging operators who benefit from marine resources represent a valuable and underexploited avenue for scientific data collection (De la Cruz-Modino & Cosentino, 2022; Martin et al., 2016, Suciu et al., 2025).

The Azores region represents an ideal setting for non-invasive sample collection through participatory science, particularly during the summer months. Approximately 90% of the samples were collected between May and September. This period coincides with the presence of sperm whales in the region, which serves as an important feeding ground, and with favorable sea surface temperatures ranging from 19 to 26°C that are known to enhance skin sloughing (Whitehead et al., 1990) in addition to the clear visibility of the sea near the surface. These conditions facilitate the detection of fecal matter and sloughed skin at the surface. This summer season also corresponds to a peak in tourism activity, characterized by a high frequency of whale-watching excursions, thereby increasing opportunities for sample collection through collaborative efforts. Such a non-invasive sampling approach could undoubtedly be successfully replicated in other regions with favorable weather conditions and active whale-watching tourism.

### 4.2. Samples predominantly reflected individual whales

Our encounters predominantly involved adult females and immature sperm whales, while mature males were observed only infrequently, consistent with previous studies (Boys et al., 2019; Steiner et al., 2012; van der Linde & Eriksson, 2020). Azorean sperm whales are numerous and highly mobile, and their core habitat is suggested to lie within Macaronesian waters (Boys et al., 2019).

Samples were collected across multiple days, over an extended period and from this large number of individuals. This studies sampling design reduced the likelihood of repeatedly sampling the same individuals. Previous studies in the region have reported low re-encounter rates (Pinela et al., 2009; van der Linde & Eriksson, 2020). Supporting this, aerial photo-identification conducted concurrently with sampling of this study, detected no recaptured individuals between 2022 and 2024 on the north coast of São Miguel, except for those made on the same day and in the same area during research dedicated fieldwork (O’Callaghan et al, 2024). Accordingly, the probability of resampling the same individual over time in our study area was very low, although particular attention was given to potential same-day, same area recaptures.

During dedicated research trips, generally only a few samples were collected per trip, typically from individuals that were photo-identified. In contrast, samples obtained through the participatory whale-watching program generally lacked photo-identification but usually consisted of a single sample per trip. Across the study period, only a small number of individuals were re-sighted: four within the same day and area, and two across different days. Notably, the mature male “Mr Liable”, identifiable by distinctive fluke and body markings, is a well-documented individual exhibiting strong site fidelity to the Azores, with records spanning from 2004 and annually since 2010 (van der Linde & Eriksson, 2020), an unusual pattern that stands out compared with other mature males (Steiner et al. 2012, Lydersen et al. 2025). Additionally, a mature female (PmM76) was re-sighted two months apart in 2024 off the north coast of São Miguel.

For re-encountered animals, multiple types of samples (skin or feces) enabled validation of the methodology. For individuals lacking fluke identification, undetected re-encounters cannot be entirely excluded; however, the probability to repeatedly sampling the same individual remained very low based on the photo-identifications made during this study.

### 4.3. Sperm whale genetic diversity in the Azores

Sperm whale have a low mitochondrial DNA diversity globally (Alexander et al., 2012). In their study encompassing Indian, Pacific and Atlantic basins, Alexander et al. (2016) described 41 MCR haplotypes. Across the North Atlantic, six MCR haplotypes (A, B, C, BB, N, X) have been determined (Alexander et al., 2016; Engelhaupt et al., 2009; Lyrholm & Gyllensten, 1998; Morin et al., 2018; Table S2), along with a unique haplotype (Y) restricted to the Gulf of Mexico (Ortega-Ortiz et al., 2012). In this study, three haplotypes (SMG_A, SMG_B, SMG_C) were identified corresponding to the three most common North Atlantic haplotypes (A, B, C) (Alexander et al., 2012). The haplotype N, previously reported in the Canaries (Engelhaupt et al., 2009), was not detected in the Azores. Importantly, we identified a new haplotype (SMG_PP), defined by a new polymorphic site, a contribution to the species genetic diversity, particularly in a basin that remains less studied compared to other locations.

In terms of genetic diversity, the observed low nucleotide diversity (π) and moderate haplotype diversity (*Hd)* determined in this study were comparable to the values obtained from the North Atlantic dataset (Table S11). They could suggest a historic population bottleneck followed by recent demographic expansion. Morin et al. (2018) attributes the primary cause of the low mitogenome diversity to a global population decline during the interglacial period. However, these results could also reflect the historical depletion of sperm whales in the Azores due to whaling followed by several decades of slow recovery (Whitehead & Shin, 2022).

Given that matrilinearity for sperm whale social groups has been demonstrated (Sarano et al., 2021), each mtDNA haplotype is expected to be found in several social units or groups that have a common female ancestor (Cantor et al., 2019). Moreover, because overall genetic diversity in the species is low, it is possible that a same mtDNA haplotype is shared among multiple social units. Individuals with a given haplotype in our samples may therefore belong to different social units, some of which have already been identified off São Miguel in van der Linde & Eriksson (2020) study. Finally, because vocal clans are suggested to be composed of social units with similar mtDNA haplotypes (Rendell et al., 2012), future studies should also incorporate acoustic data.

On one occasion, two different individuals diving together off the south coast of São Miguel, revealed distinct mitochondrial haplotypes (PmM13 possessed the haplotype SMG_A and PmM14 the haplotype SMG_B; overlapping points on Figure 3). Although sperm whales are known to exhibit strong matrilineal social structure (Whitehead, 2003), associations among genetically unrelated individuals have also been reported (Christal et al., 1998; Konrad et al., 2018; Sarano et al., 2021).

### 4.4. Haplotype differential distribution around São Miguel

Despite the large area covered by the sperm whales in Macaronesia, our results reveal a clear segregation of sperm whale individuals correlated to their mtDNA haplotypes around São Miguel: individuals carrying the haplotype SMG_C tend to cluster on the south of the island, which clearly distinguished them from the other haplotypes.

This fine-scale geographical structure in a highly mobile cetacean supports the notion that sperm whales show site preference (Hoelzel et al., 2009; Lyrholm & Gyllensten, 1998), as it has been documented among vocal clan in the Lesser Antilles (Vachon et al., 2022). Our findings extend this concept by suggesting that such spatial structuring can occur at an even smaller scale, here around a single island, such as São Miguel.

We hypothesize that different matrilineal lineages may preferentially occupy different habitats, which has important implications for understanding sperm whale ecology and for conservation. Our data support the idea that individuals travel to specific sites characterized by geography or habitat features, where knowledge of feeding areas can be socially transmitted (cultural ecological specialization) (Vachon et al., 2022). The observed site fidelity may be advantageous for hunting, as familiarity with local prey patches, particularly cephalopods, could improve foraging success and reduce energic costs. Consistent with this, individuals of different haplotypes showed variation in the mean ocean depth of their dives, a pattern also reported among social structures on the west coast of Martinique (Laurent et al., 2025) and consistent with models predicting spatial variation according to bathymetry among different social groups (Pace et al., 2018).

Territoriality cannot be ruled out either, where individuals would avoid each other and/or preferentially associate with kin. Such behavior could explain the preferential distribution of the SMG_C haplotype, but it could also happen on the North of São Miguel where multiple haplotypes co-occur. In one documented instance of agonistic behavior (head butting) on the 23th of August 2022, skin fragments obtained from two young males revealed different haplotypes (PmM15: SMG_B and PmM16: SMG_PP) (Burslem et al., 2026).

These findings have direct conservation implications. The distinct haplotype distribution between the north and south coasts of São Miguel suggests that these areas harbor different matrilineal lineages and should be managed as separate units. Anthropogenic pressures such as whale-watching activity, vessel traffic, and noise disturbance may therefore affect genetically and socially distinct groups differentially, warranting site-specific management measures. Future research should expand both sample sizes and geographic coverage to the other islands of the archipelago, in order to fully characterize the genetic diversity of Azorean sperm whales and identify additional fine-scale structuring that may have conservation relevance.

### 4.5. Sperm whale mature male site fidelity

Knowledge about the site fidelity of mature male sperm whales and the relationship with their mitochondrial haplotypes and those of matrilineal groups remains limited. In the Indian Ocean, males sampled in Mauritius were genetically unrelated to females visited in Mauritius (Girardet et al., 2022). In our study, the mature males shared common haplotypes with the females and juveniles. Specifically, Mr Liable possesses a common haplotype with the local females (SMG_C), the most common haplotype in the south of São Miguel, where this mature male was sighted several times over the course of this study. Similarly, the other mature males of this study, PmM49 (SMG_C) and PmM83 (SMG_A), also shared common haplotypes with the females and juveniles. This pattern might be due to the low genetic diversity in the North Atlantic, where the haplotypes in question are among the most frequent, but also due to the limited number of samples from mature males. Further investigation of haplotypes in mature male sperm whales, including individuals from higher latitude such as males recorded in Arctic Norway (Morange et al. 2025) that are known to migrate to the Azores (Steiner et al. 2012; Lydersen et al., 2025), could help determine their genetic lineages.

## 5. Conclusion

This study highlighted the effectiveness of non-invasive genetic sampling, such as sloughed skin, and feces, in the conservation genetics of sperm whales. Notably, it was the first to use fecal samples to establish sperm whale haplotypes, and it allowed to identify a new haplotype for the species. The findings from the haplotype distribution around a relatively small area (São Miguel Island) contribute to a better understanding of the population sub-structure of sperm whales in the Azores. It also shows the significance of matrilineal local groups and their possible small scale site preferences when evaluating population structure. Sperm whales have a complex social structure that leads to geographical and habitat preferences. Their distribution around São Miguel Island was not homogenous and described patterns that may inform predictive trends, which need to be considered in conservation management and sustainable tourism practices.

## Author contributions

Stéphanie R.A. Suciu: conceptualization, data curation, methodology, investigation, visualization, writing – original draft, writing – review and editing, funding acquisition, project administration. José M.N. Azevedo: resources, writing – review and editing, funding acquisition, project administration, supervision. Sean O’Callaghan: data curation, visualization, writing – review and editing. Bruno Serranito: formal analysis, writing – review and editing. Jean-Luc Jung: conceptualization, resources, validation, writing – review and editing, supervision.

## Acknowledgments

This work originated as part of Stéphanie Suciu’s PhD thesis “Ecology of cephalopod/cetacean interactions in the Azores” at the University of the Azores (Suciu, 2025). Permission to approach cetaceans and collect non-invasive samples was granted by the relevant authorities under permits AMP/2020/014, AMP/2022/007, DRAM/LEMASM/2023/008, and DRAM/LEMASM/2024/008 (Direção Regional dos Assuntos do Mar); 20/2022/DRCTD (Direção Regional da Ciência e Transição Digital); LMAS-DRPM/2023/07 and LMAS-DRPM/2024/10 (Direção Regional de Políticas Marítimas); and CCIR-RAA/2023/36 and CCIR-RAA/2024/45 (Direção Regional da Ciência, Inovação e Desenvolvimento). We thank the dedicated whale-watching community for their contributions to the collection of cetacean biological samples, including CW Azores, Dive Azores, Espaço Talassa, Futurismo, Norberto, Sea Colors, and Terra Azul. Miguel Carvinho (Terra Azul) also provided essential operational support during the dedicated research trips off the south coast of São Miguel: thank you to Stephanie Almeida, Jorge Armal, Sanne Bakker, Tiago Batista, Marylou Féat, Filipe Ferreira, Sandra Gonzalez, Nicole Pereira, Nuno Pimentel for their support during the 2023 and 2024 field seasons. Thank you to Rui Rodrigues (Futurismo), as well as Rafael Martins, Anaïs Builly, Pablo Varona, and Vítor Silva for their support to collect Mr Liable samples in 2023. Thanks are extended to Tomás Anselmo and Paulo Luís Sousa for facilitating the fieldwork on the north coast of São Miguel (2022-2023). We are grateful to Francisco Garcia, Inês Coelho and Inês Subtil for their support throughout this work. We thank João Faria, Duarte Toubarro, Rachel Haderlé, and Karen Cosnier for their support in the laboratory work.

